# The relationship between the waiting time of postoperative radiotherapy and the prognosis of high grade glioma:a systemic review and meta-analysis

**DOI:** 10.1101/2020.01.10.901447

**Authors:** Guang-lie Li, Shuang Lv, Ying Xu, Hai-bo Zhang, Ying Yan

**Author notes:** Corresponding author, Department of Radiation Oncology, General Hospital of Northern Theater Command, Shenyang city, China. First author (Guang-lie Li). These authors also contributed equally to this work (Shuang Lv, Ying Xu, Hai-bo Zhang).

## Abstract

Objective: The relationship between the waiting time of postoperative radiotherapy and the prognosis of patients with high-grade glioma is still inconclusive, and we addressed this issue through a systematic review and meta-analysis. Methods: Twenty studies published between 1975 and 2019 about waiting times (WT) of radiotherapy with high-grade glioma were retrieved for meta-analysis.The meta-analysis was performed by converting the effect sizes of different WT into regression coefficients (β) and standard error (SE) to indicate the daily impact of delay on OS. Results: A total of 8462 high-grade glioma patients were included in the 20 studies, and no correlation between WT delay and OS was found in the unadjusted model through meta-analysis (HR=1, 95%CI=0.99-1.01, *p*=0.962). Meta-regression was used to adjust for other prognostic factors and no clear evidence of the relationship between WT delay and OS was found. Conclusion: This meta-analysis suggests that there is no clear evidence for the effect of delayed radiotherapy on OS with high-grade glioma patients.

## 1. Introduction

High-grade gliomas are originated from glial cells and are common primary intracranial tumors in adults.According to the World Health Organization (WHO) classification, high-grade gliomas mainly include grade III anaplastic glioma and grade IV glioblastoma multiforme.Compared with low-grade glioma, high-grade glioma grows faster and is more invasive.Its predilection age is 50-60 years^[1]^.It also has the characteristics of high morbidity, high recurrence rate and high fatality rate.Currently, the major treatment for high-grade glioblastoma includes surgery, chemotherapy and radiotherapy. Despite the positively comprehensive treatment, the prognosis of patients is still poor. According to the data in previous years, the 2-year survival rate of glioblastoma patients is only 27%, while only 10% of patients survive for more than 5 years^[2]^.

Because high-grade glioma is a diffuse and invasive intracranial malignant tumor, it is still difficult to completely remove the tumor even with advanced techniques such as intraoperative image guidance^[3]^.Therefore, the American NCCN guidelines^[4]^ and the European ESMO guidelines^[5]^ suggest adjuvant radiotherapy after surgery, concurrent chemotherapy on the first day of radiotherapy, and continued chemotherapy after radiotherapy.In previous studies, the relationship between delayed radiotherapy and prognosis has been explored in various types of tumors (such as breast cancer, head and neck tumors)^[6-7]^.Clinical and experimental data support the idea that delayed radiotherapy may reduce the local control rate of the tumor and thus affect the overall survival of the patients.However, the waiting time of radiotherapy for high-grade gliomas is still controversial.Several studies have shown unexpected survival advantages for groups with longer waiting times, and there is no clear radiobiological explanation.Ultimately, it remains to be seen whether delayed radiotherapy affects survival of patients with high-grade glioma, so we propose a systematic review and meta-analysis to address about the impact of delayed radiotherapy on survival of patients with high-grade glioma.

## 2. Materials and Methods

### 2.1. Search strategy

Our goal was to identify all published studies of postoperative delayed radiotherapy with high-grade glioma patients, regardless of study design, language and format. The retrieval date was between January 1, 1975 and November 1, 2019.For published articles related to the topic were continuously monitored until finalization.Pubmed, EMBASE, Cochrane were used for retrieval.Search keywords include:watiting, time, timing, interval, delay, prolong, initiation, glioma, glioblastoma, radiation, radiotherapy, chemoradiation, chemoradiotherapy, adjuvant(see Appendix 1 for the detailed retrieval scheme in Supplementary data).If research reports provided incomplete information or vague descriptions, we tried to contact the authors for more information.

### 2.2. Inclusion and exclusion criteria

Eligible studies were need to meet the following criteria:neurosurgery (biopsy, partial resection or total resection), pathologic diagnosis of high-grade glioma (grade III or IV), adult, postoperative radiotherapy, accepted concurrent chemotherapy or not, regardless of radiotherapy course and radiotherapy dose.Additional inclusion include:(1) OS was one of the results of the study; (2) the time from surgery to radiotherapy was defined as the time interval between the first neurosurgical operation and the beginning of radiotherapy, and the results were described as continuous variables or category variables; (3) the clinical and therapeutic characteristics and outcomes of patients with high-grade glioma (grade III/IV) were clearly reported.

Exclusion criteria include:(1) patients with recurrent high-grade glioma; (2) patients with low-grade glioma; (3) received chemotherapy or other adjuvant therapy before the beginning of radiotherapy; (4) non-human experimental studies; (5) OS was not reported according to the interval between surgery and radiotherapy. Study design, language and publication format were not limited.

### 2.3. Data extraction

We extracted the following data: study author, publication date, study period, number of patients, adverse prognostic factors (KPS, age, surgical resection), chemotherapy, radiation dose, waiting time (WT) and subgroup,effect of delayed radiotherapy on OS, publication format.No more information was obtained by contacting the author. In order to perform the meta-analysis, the study must have the hazard ratio(HR) and the confidence interval (CI) or median survival for the WT subgroup.

In order to synthesize the study results of different WT variables (such as continuous variables and category variables), we converted the effect size of WT for each study into regression coefficient(β) and standard error(SE) to represent the impact of WT on OS.For studies with categorical WT, the median of each group was used to represent the center value of each WT category.For the closed interval, took the midpoint of the two intervals, and for the open upper interval, took 1.2 times for the minimum value to represent its size^[8]^.The regression coefficients(β) and standard error(SE) conversions for continuous, dichotomous and ordinal data were as follows:(1)For continuous variable of WT, the β was calculated as log (HR), and the SE was calculated as (log (upper CI) −log (lower CI)) /3.92^[6]^. The unit of HR was converted into days before calculation.(2)For studies with only 2 WT groups,the β calculated as log(HR)/([x_n_-x_0_]*3.92), and the corresponding SE calculated as (log(upper CI)-log(lower CI) /([x_n_-x_0_]*3.92), where CI was the confidence interval, x_n_ represented the exposure level of the n group, and x_0_ represented the exposure level of the reference group^[9]^.If only HR and *p* values were provided, upper CI and lower CI were calculated according to the formula proposed by Altman^[10]^, and then substituted into the above formula.(3)For two or more groups of studies, if the total number of people and the number of events reported in the study,then GLS(Generalized least square) model was used to estimate the β and SE, if not reported, VWLS (variance-weighed linear square) model was used to estimate the β and SE^[11]^.

### 2.4. Meta-analysis procedures

The fixed effect model was used to combine the adjusted regression coefficients of each study.The inverse variance (1/SE^2^) was used to weigh individual studies.For the consistency evaluation of all included studies, conventional statistical tests (Cochran’s Q) were first used for evaluation, and I^2^ was further used.If I^2^ was greater than 50%, indicating greater heterogeneity^[12]^.In the next step, the funnel plot was drawn. The evaluation of selection bias was evaluated by Begg and Egger’s test^[13]^.The symmetrical inverted funnel shape, *p*>0.05 was generated from well-behaved data sets. Ultimately, it showed that publication bias was impossible.Meta-regression was used to adjust for other prognostic factors(KPS,age,surgical resection) to clarify the relationship between the waiting time and the prognosis of high-grade glioma.Statistical software STATA 15.0 version was used for statistical analysis.

## 3. Results

A total of 11186 references were retrieved in different databases, and 158 references were retrived for full-text. Excluding referrences which duplicated or can’t find the full-text or abstracts, only 44 articles were examined for full-text.After excluding unqualified studies according to the inclusion criteria, 27 studies were included for qualitative synthesis.Finally,20 studies were chosen for Meta-analysis(Figure 1).The studies were published between 2007 and 2019, and 18 of them had full texts available, while two acquired data only from abstracts.There were 8462 patients with high-grade glioma, with sample sizes ranging from 50 to 2,535(Table 1).Four of the studies were expressed as continuous data, eleven as two group, and five as three group.One study was analyzed by using the NRG tumor /RTOG database, and one study analyzed the relationship between delayed radiotherapy and the prognosis of high-grade glioma based on MGMT promoter methylation status. With the exception of two studies, almost all of them focused on the relationship between WT and OS.

**Figure 1.**
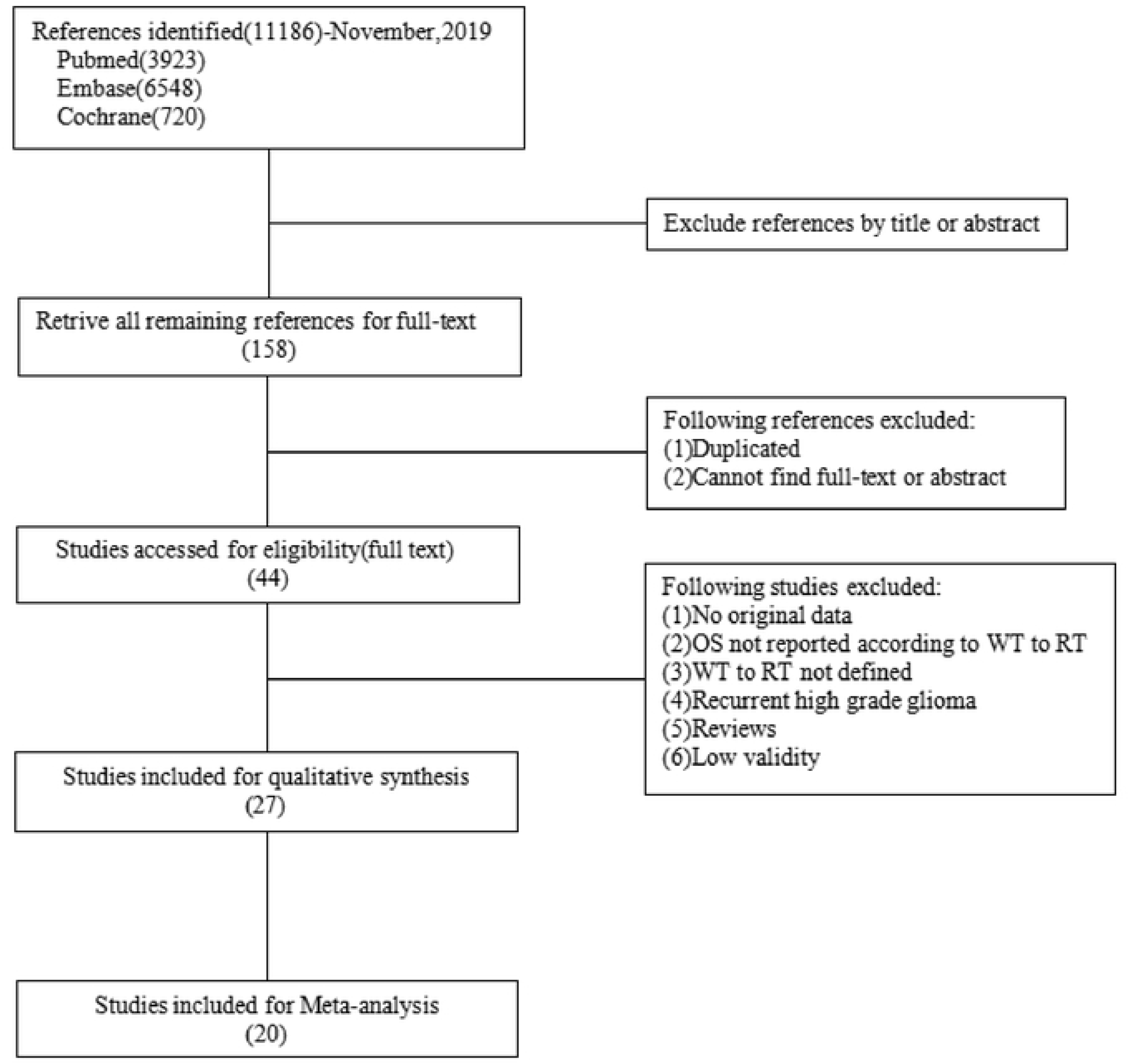
Flow chart of the study selection

**Table 1.**
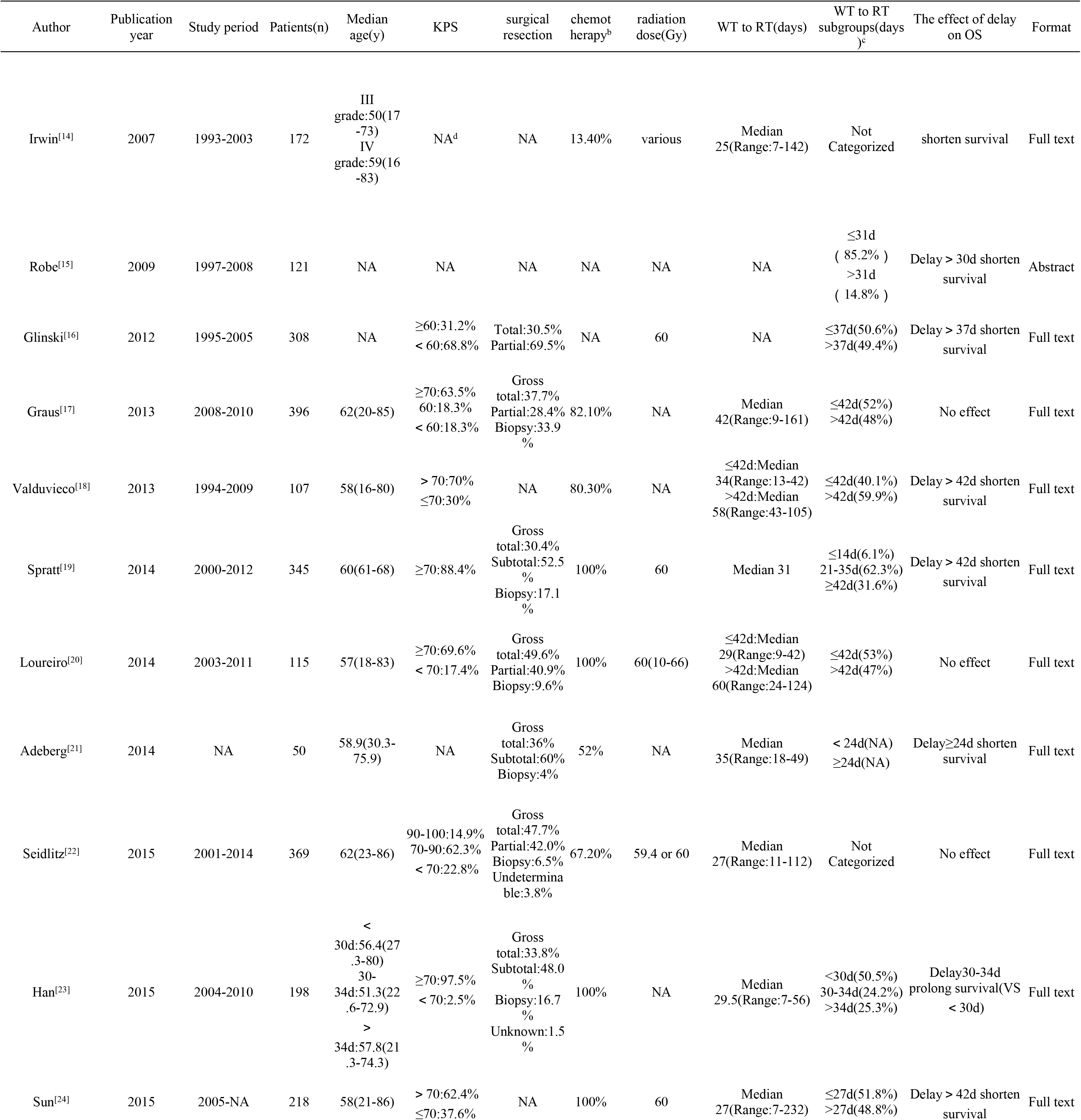
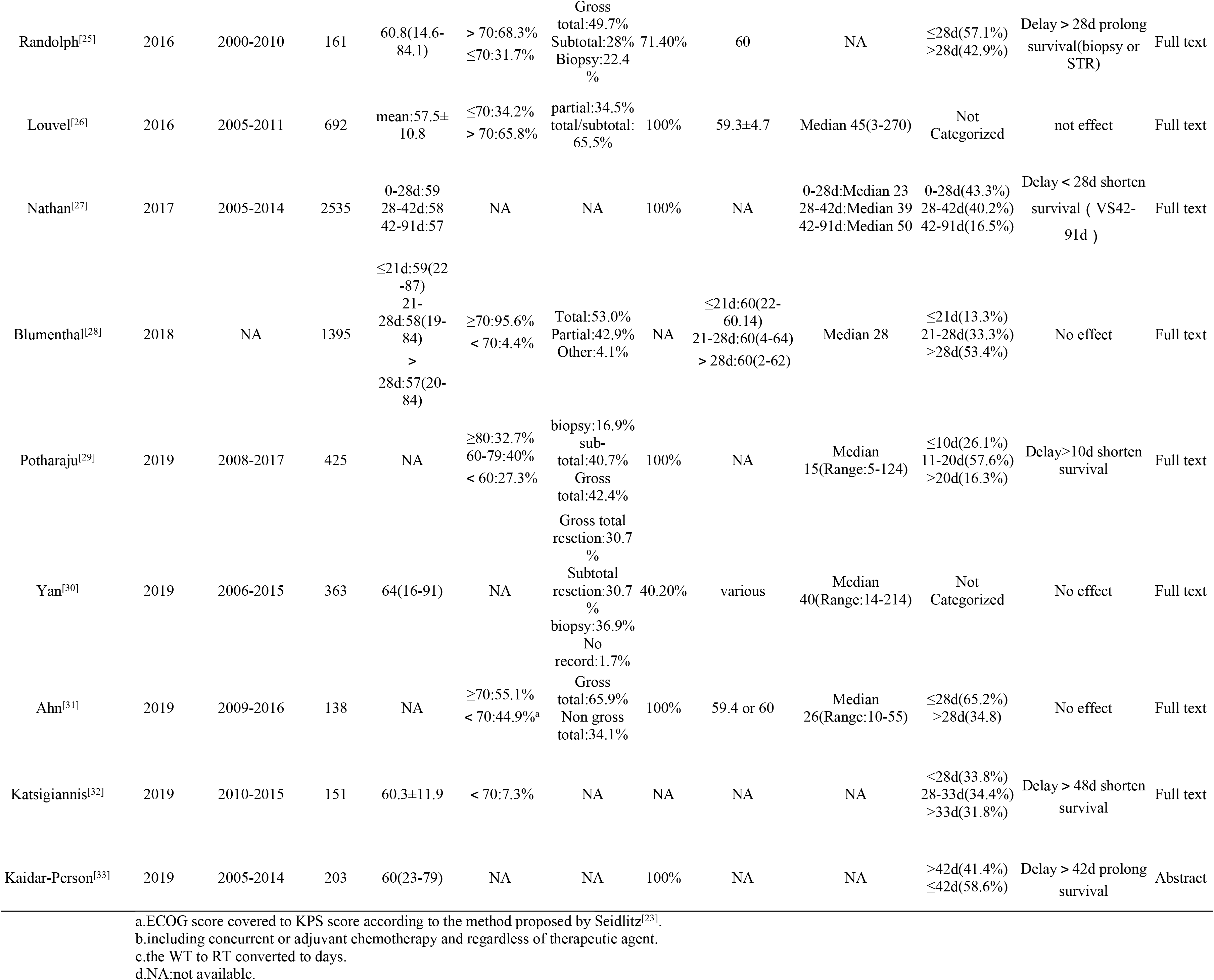
Characteristics of studies

The effect of delayed radiotherapy on patients with high-grade glioma had conflicting results in the 20 studies.Among them, 7 studies reported that delayed radiotherapy had no significant relationship with the prognosis of patients with high-grade glioma, while 9 studies reported that delayed radiotherapy significantly shorten the survival of patients with high-grade glioma, but each study set a different cut-off value.One suggested that early radiotherapy was beneficial to high-grade glioma patients whose MGMT were methylated among 9 studies.However, four studies reported additional survival benefits from delayed radiotherapy in patients with highgrade glioma.One reported that in the case of biopsy or partial resection, delayed radiotherapy was more conducive to survival of patients with high-grade glioma. One suggested that moderately delayed radiotherapy (30-34d) prolonged survival of patients, while the other two studies suggested that delay> 42d was more suitable for long-term survival of patients.HR values for different groups of each study are listed in Table 2.

**Table 2.**
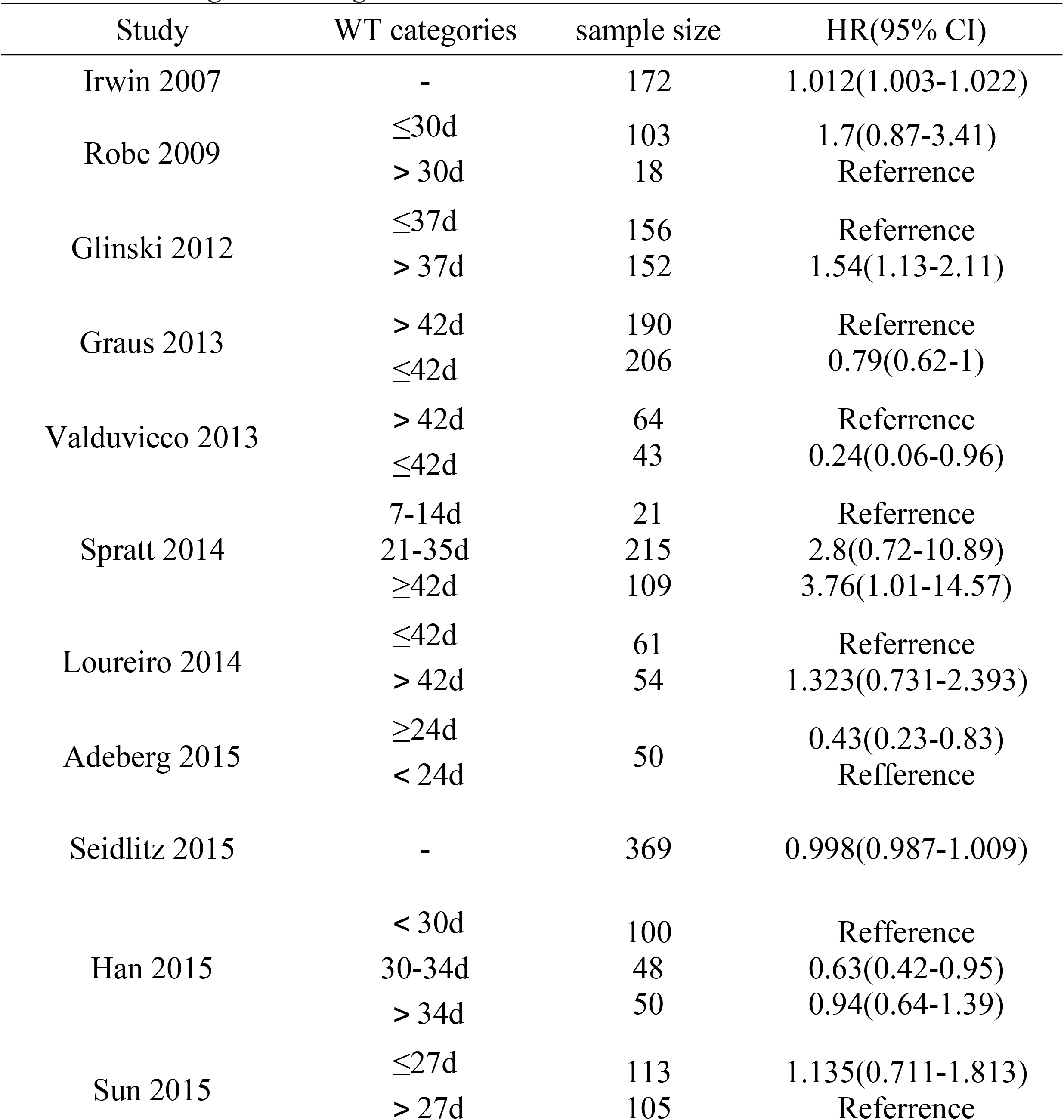
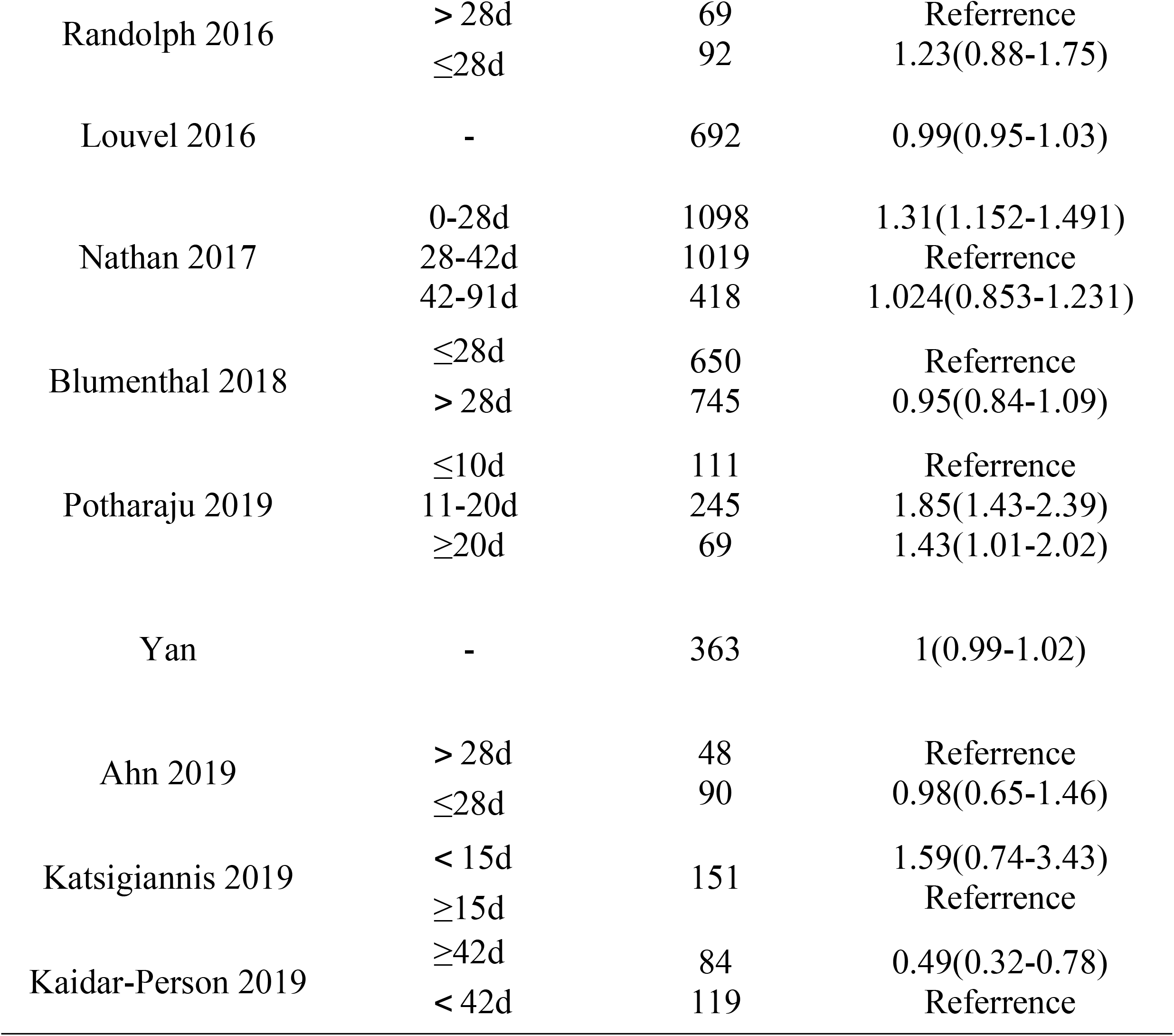
Waiting time categories and clinical outcomes

To solve this problem, we combined the entire cohort for analysis and found no evidence of the relationship between delayed radiotherapy and OS (HR= 1, 95%CI=0.99-1.01, *p*=0.962)(Figure 2).No evidence of significant statistical heterogeneity was found in these studies (Cochrane’s Q test *p* > 0.05, I^2^=26.7% < 50%).The funnel plot was drawn to show a symmetrical design. The regression test of Begg (*p*=0.770) and Egger (*p*=0.719) showed no obvious asymmetry, indicating no clear evidence of publication bias(Figure 3,4).The prognostic factors were adjusted by meta-regression, there is still no clear evidence that the waiting time of radiotherapy has an impact on the prognosis of high-grade glioma patients (Table 3).

**Figure 2.**
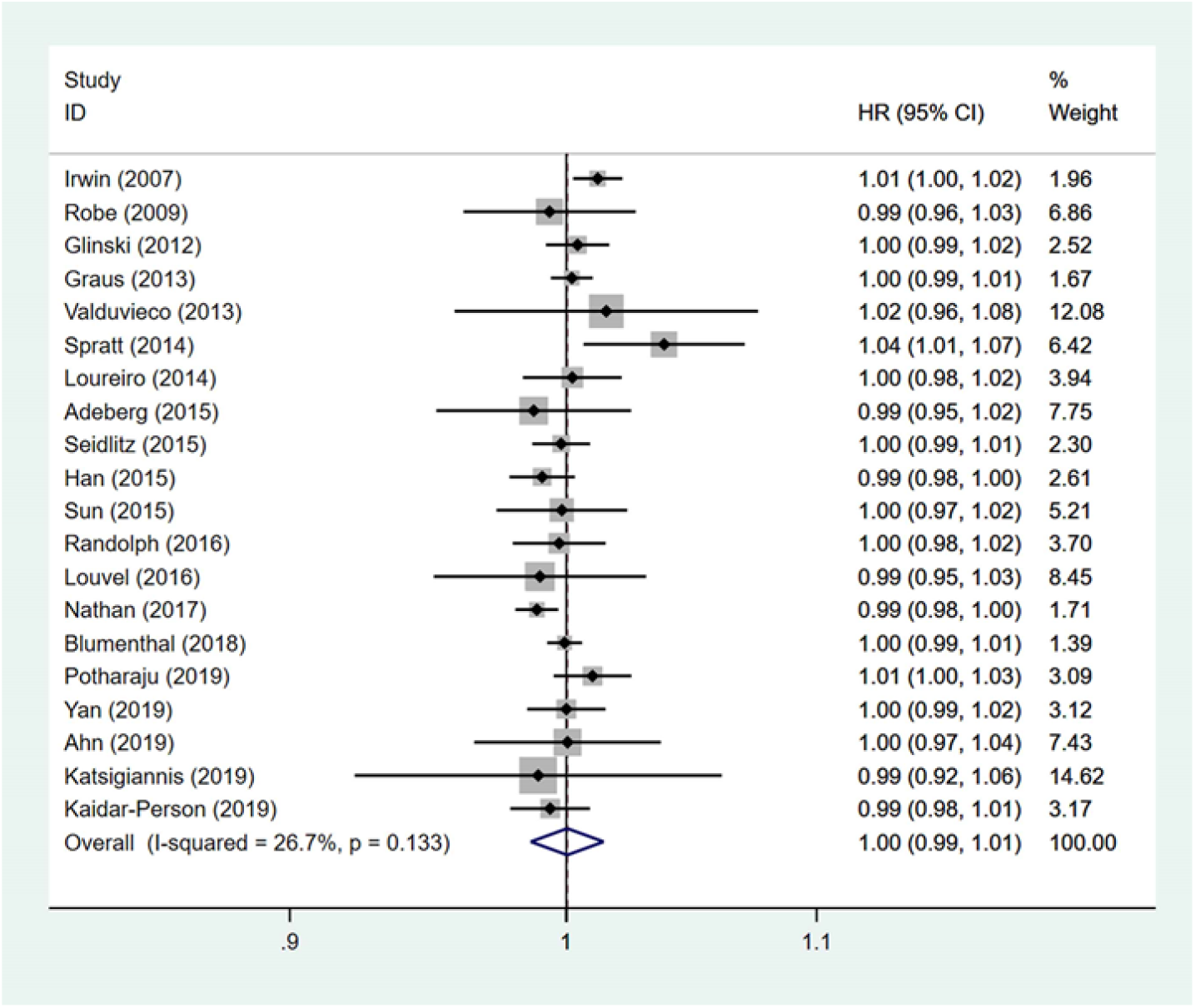
Forest plot.

Shows each study and overall hazard ratio(HR) per day of delay with 95% confidence interval(CI) for OS.The square size is proportional to the weight assigned to the each study based on 1/SE^2^.For the combined results, the length of the diamond represents the 95% CI of the summary.

**Figure 3.**
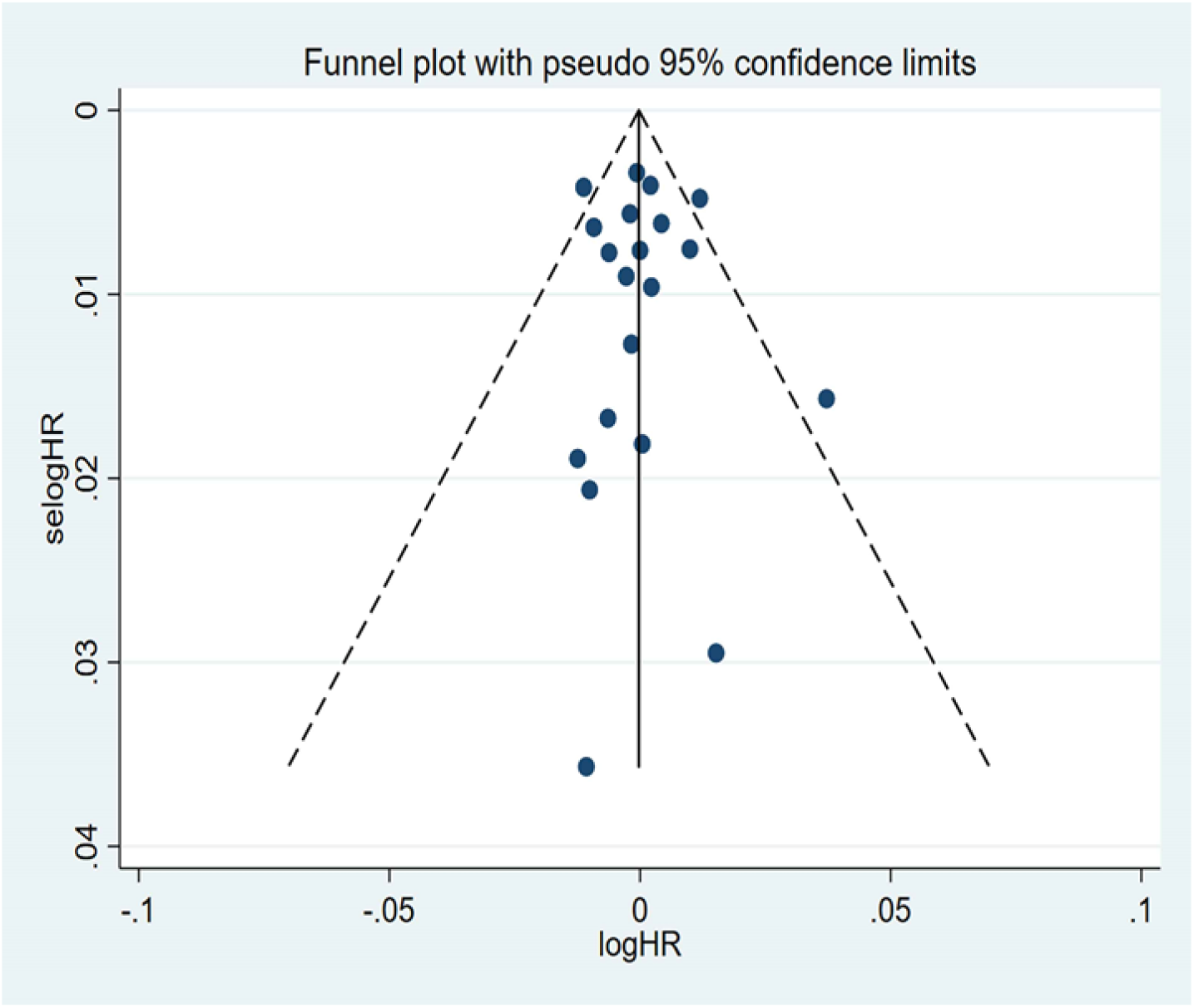
Funnel plot.

Shows the relationship between the log harzard ratio and standard error of the log harzard ratio.The harzard ratios(HR) estimated are the effect of waiting time for every day.The dotted line represents the combined HR for all OS studies.The filled circle represents 20 studies that considered potential publication bias.

**Figure 4.**
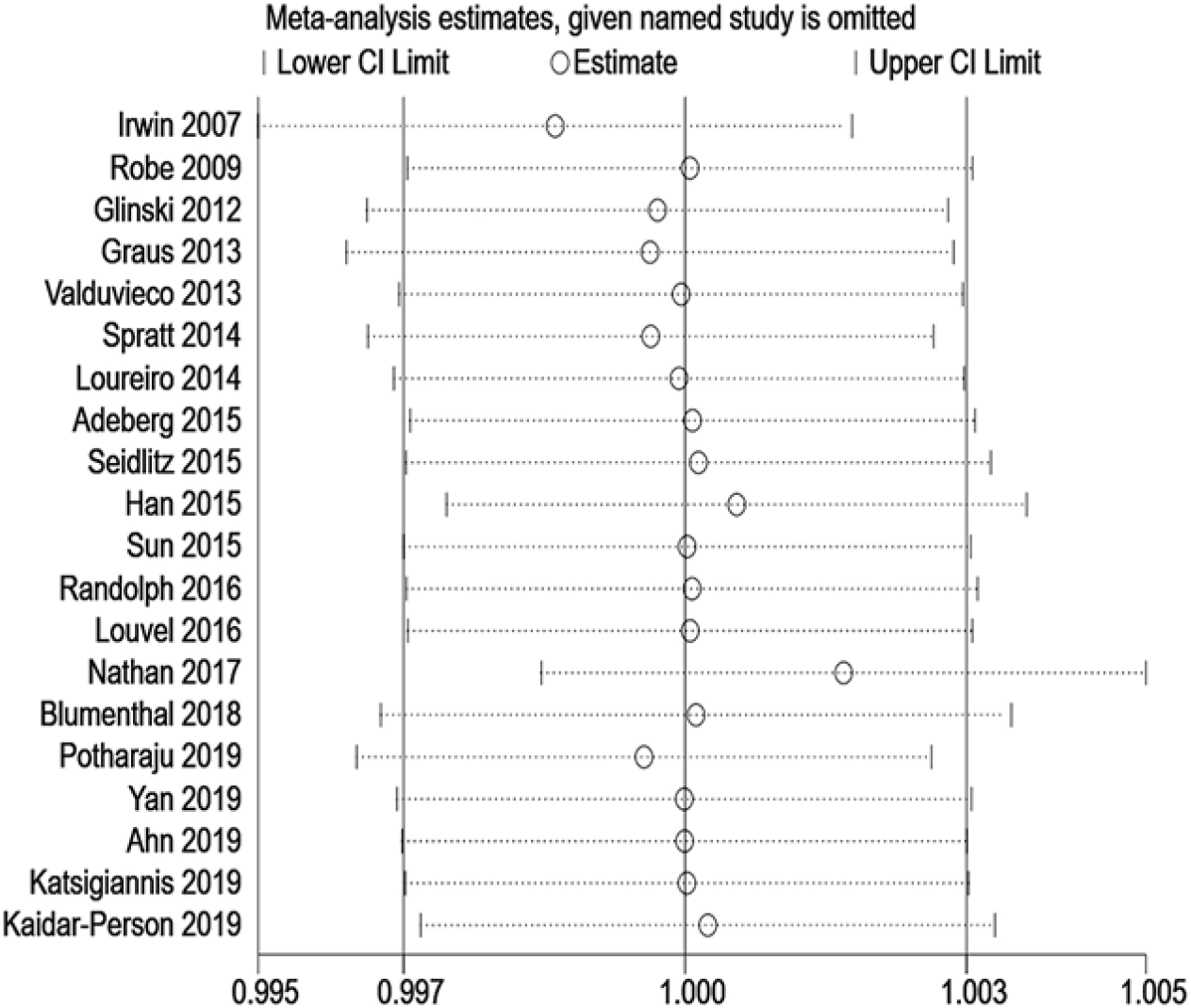
Sensitivity analysis.

Shows the impact of individual studies on the pooled HR for OS with high grade glioma. The vertical axis indicates the overall HR and the two vertical axes indicate its 95% CI. Every hollow round shape indicates the pooled HR when the left study is omitted in this meta-analysis. The two ends of every broken line represent the respective 95% CI.

**Table 3.** Meta-regression adjusted for prognostic factors

| Prognostic factors | HR | 95%CI | *p* value |
| --- | --- | --- | --- |
| Age<sup>14,16,19-22,24-27,30-32</sup> | 1.002 | 0.990-1.014 | 0.708 |
| KPS<sup>a</sup> < 70%<sup>14,18-22,24-26,31-32</sup> | 1.002 | 0.998-1.006 | 0.321 |
| Surgical resection |  |  |  |
| gross total or subtotal<sup>14,17,19-22,25-26,30-31</sup> | 0.994 | 0.980-1.007 | 0.307 |
| biopsy<sup>14,20,25,30</sup> | 1.000 | 0.976-1.025 | 0.934 |

a. Adeberg<sup>[21]</sup> reported HR of KPS>70% and it was converted according to the method proposed by Tierney<sup>[34]</sup>.

## 4. Discussion

High-grade glioma is the most common malignant primary brain tumor in adults, and despite multimodality therapy such as surgery, chemotherapy and radiation, the median overall survival is still short, for 16 to 20 months^[35]^.At present, there is still a great dispute about when to start radiotherapy for high-grade glioma, and there is no unified conclusion.Since Do^[36]^ first reported the effect of waiting time on the prognosis of patients with high-grade glioma, more and more studies have been trying to solve this problem in the past 20 years,but there is still no clear answer.

Someone who advocate early radiotherapy believe that high-grade glioma has a fast growth rate and a short doubling time, about 38-51h^[37-38]^.Therefore, delaying radiotherapy is not conducive to the prognosis of patients. Moreover, rapidly proliferating tumor cells are more sensitive to radiation than slowly proliferating tumor cells^[39]^.Some scholars proposed the Gompertz equation, which combines the concept that larger tumor cells grow more slowly^[40]^.The cell growth described in this equation showed exponential properties in the early stage, then gradually reached saturation and approached the plateau stage with the increase of tumor size^[41]^.When the tumor is surgically removed, the residual small tumor cells grow more rapidly^[41]^, making them more sensitive to radiation.These explanations seem to make clear conclusions.

However, some researchers hold the opposite opinion, believing that too early (within 3 weeks) radiation treatment causes more serious neurological damage^[42]^. Moreover, local hypoxic cells increased in the early postoperative period are insensitive to radiation^[43]^, which affect the efficacy of radiotherapy.These hypoxic areas are reoxygenated when the wound healed and blood was re-supplied.Therefore, postoperative radiotherapy should be postponed until the postoperative hypoxia disappeared.It could avoid hypoxia-induced radiation resistance and obtain the optimal activities for killing tumor, but the optimal time has not been determined^[44]^.

At the beginning of this topic,after a comprehensive analysis of 4 studies,Lawrence^[44]^ suggested that moderate delayed waiting time (no more than 4-6 weeks after surgery) is safe and may have some benefits.In contrast, there was no evidence that waiting times for more than six weeks are reasonable.However, through further meta-analysis for 12 studies, Loureiro^[45]^ found no clear correlation between radiotherapy delay and the prognosis of patients(HR=0.98, 95%CI =0.90-1.08, *p*=0.70).Moreover, there are still many studies going on in recent yeears and reporting different results.Therefore, it remains a mystery whether radiotherapy waiting time affects the prognosis of patients with high-grade glioma.

In our meta-analysis, we added the latest results in recent years to find the relationship between delayed radiotherapy and the prognosis of patients with high-grade glioma, and found different conclusions from these 20 studies.Seven studies included more than 3400 patients, each of which showed no effect of delayed radiotherapy on OS.Nine studies included more than 1,900 patients, reporting that prolonged radiotherapy shortened OS, but not all of them set the same cut off value.Three of the studies suggested that delayed radiotherapy more than 42 days were significantly detrimental for long-term survival.Finally, four studies involving more than 3,000 patients reported additional survival benefits from prolonged radiotherapy.One study suggested that mildly prolonged (30-34d) radiotherapy was suitable for long-term survival of patients, one study suggested that delayed radiotherapy improved survival in patients who underwent biopsy or partial resection.Fuhtermore, one study reported that radiotherapy initiated longer than 42 days prolonged survival of patients.This meta-analysis was based on data from 20 observational studies, including 8462 patients with high-grade glioma,.There was no evidence that delayed RT affected OS (HR= 1,95%CI=0.99-1.01,*p*=0.962).However, in clinical practice, the prognosis of patients with high-grade glioma is affected by many factors (such as age, KPS score, surgical resection). After adjusting these confounding factors, there was still no clear evidence that the waiting time of radiotherapy affected the prognosis of patients with high-grade glioma. These results are basically consistent with those obtained from Loureiro^[45]^.

Theoretically, there is no suitable reason to explain the delay of radiotherapy, except that patients with postoperative incision infection need to be treated for a period of time or the patients are in poor physical conditions.It seems unnecessary to discuss the topic.However, in clinical practice, it has been found that many factors, including the balance between supply and demand of radiotherapy, the popularity of radiotherapy in some areas, the economic and social status of patients and their families, as well as their education level, can affect the time of starting radiotherapy^[46]^. And the delayed radiotherapy has shown additional survival benefits in some studies.

The most representative article published by Nathan^[27]^,which included 2,535 patients with high-grade glioma and was divided into three groups according to different time intervals (0-28d, 28-42d, 42-91d).It was found that patients who started radiotherapy at 42d after surgery had a better prognosis than those who started radiotherapy within 28d after surgery.Kaidar-person^[33]^ also found that patients lived longer who started radiotherapy more than 42d.These results are conflicted with the conclusions from the early comprehensive analysis by Lauwrence (that the prognosis of patients is poor while radiotherapy delayed over 6 weeks). However, no radiobiological results have been reported to explain this phenomenon.

In summary, our results show no significant correlation between postoperative waiting time of radiotherapy and the survival of patients, and there are some limitations in our study. First, all the studies in this meta-analysis are non-random and retrospective. Second, the HRs of waiting times we used in the analysis are not adjusted for prognostic factors in some studies. Thirdly, in the transformation of regression coefficients, an estimation method is adopted for more than two groups of data, and errors may exist in the results. Although there are some limitations in this analysis, it gives a more reliable result than a single observational study, the answer is between favorable and unfavorable.To get a more accurate answer, we may need to conduct some cell biology or animal experiments to draw a conclusion.

## Acknowledgements

None.

## Supplementary Data

See Appendix 1.

